# scIsoAgent enables autonomous isoform-resolved characterization and sequence-informed interpretation of long-read single-cell transcriptomes

**DOI:** 10.64898/2026.06.11.731519

**Authors:** Chengzhi Zhao, Mingyu Liu, Xue Li, Dongsheng Li, Yichi Xu, Zilong Wang

## Abstract

Alternative isoform usage can alter gene function independently of total gene expression, creating a need to resolve transcript isoforms at single-cell resolution. Long-read single-cell RNA sequencing meets this need by linking cellular identity to transcript isoforms and sequence-level features. Realizing its full biological value requires reproducible workflows that connect specialized long-read analysis with biological interpretation. Existing large language model (LLM)-based biomedical agents support general omics analysis, but are not designed for isoform-resolved long-read single-cell workflows. Here, we present scIsoAgent, an autonomous LLM-powered scientific agent for long-read single-cell RNA-seq analysis. scIsoAgent turns heterogeneous long-read single-cell inputs into traceable isoform-resolved workflows, using stage-aware planning and persistent computational context to support both execution and interpretation. Across complementary evaluations, this design improved the continuity from analysis planning to executable, interactive workflows compared with general-purpose LLM baselines. In real-data reanalysis, scIsoAgent recovered major findings from published long-read single-cell resources and extended a representative differential transcript usage event into a sequence-informed functional hypothesis. By linking full-length isoform sequences with model-inferred transcript properties, scIsoAgent connects observed isoform usage with potential sequence-level functional consequences. These results demonstrate that autonomous scientific agents can transform fragmented long-read single-cell analysis into coherent, reproducible workflows for isoform-resolved discovery and biological interpretation.

## 1 Introduction

Single-cell RNA sequencing (scRNA-seq) has transformed transcriptomic studies by enabling gene expression profiling at single-cell resolution. By resolving cellular heterogeneity that is masked in bulk measurements, scRNA-seq has provided important insights into cell types, cell states, developmental trajectories, disease-associated programs, and tissue-specific regulatory processes [1, 2, 3, 4]. The scalability of droplet-based platforms has further established scRNA-seq as a central approach for constructing cellular atlases and studying complex biological systems [5, 6, 7].

Despite these advances, most conventional scRNA-seq methods rely on short-read sequencing and are primarily used for gene-level expression analysis. Because many protocols capture short fragments from the 3’ or 5’ ends of transcripts, they usually cannot reconstruct full-length transcript structures or reliably distinguish isoforms. This limits the study of alternative splicing and isoform-specific regulation at single-cell resolution [8]. The limitation is biologically important because alternative isoform usage can alter gene function independently of total gene expression, changing the regulatory output of a gene in ways that gene-level measurements cannot capture [9, 10].

Long-read RNA sequencing technologies, including Oxford Nanopore Technologies (ONT) and Pacific Biosciences (PacBio), provide an effective solution by generating reads that can span complete or near-complete transcripts [11, 12, 13]. When combined with single-cell barcoding strategies, long-read sequencing enables isoform-resolved transcriptomic profiling at cellular resolution, allowing cell identity to be linked with isoform structures [14, 15]. This capability enables transcript diversity to be interpreted in its cellular context and preserves isoform-level information that is lost in gene-level abundance measurements. Consistent with this potential, our PubMed-based survey shows that long-read single-cell transcriptomics has expanded rapidly in recent years, reflecting growing adoption of isoform-resolved cellular profiling (Supplementary Fig. S1a).

This rapid growth has also been accompanied by increasing computational diversity (Supplementary Fig. S1b,c). Specialized tools have been developed for different steps of long-read transcriptomic analysis, from read processing and isoform reconstruction to transcript usage estimation and downstream interpretation [11]. Representative tools support splice-aware alignment with uLTRA [16], isoform quantification with StringTie2 [17], Mandalorion [18], TALON [19], FLAMES [14], IsoTools [20], and LIQA [21], as well as downstream interpretation and visualization with Swan [22]. Despite this expanding ecosystem, most tools address specific analytical steps rather than end-to-end isoform-resolved single-cell workflows. Such workflows are context-dependent, varying with sequencing platform, available input resources, biological objective, and upstream choices in isoform discovery or quantification. This dependence limits rigid pipelines and motivates adaptive workflow systems that can preserve reproducibility across heterogeneous long-read single-cell analyses.

Recent advances in large language models and AI agents have created new opportunities for biomedical data analysis and scientific workflow assistance. Existing systems have explored agentic support for single-cell analysis [23, 24, 25] and broader biomedical research workflows [26, 27, 28]. However, these systems are not designed for the specific computational structure of long-read single-cell transcriptomics, where isoform-resolved analysis requires coordinated handling of platform-specific inputs, transcript-level evidence, cell-level context, and sequence-informed interpretation.

To address this need, we developed scIsoAgent, an autonomous LLM-powered agent for long-read single-cell RNA-seq analysis. scIsoAgent provides an adaptive workflow framework for ONT and PacBio-based data, connecting heterogeneous long-read single-cell inputs with isoform-level analysis and sequence-informed biological interpretation. By combining stage-aware planning, traceable script-based execution, and persistent computational context, scIsoAgent supports reproducible analysis across interactive workflow steps. This design reduces the practical barriers to isoform-level discovery in single cells and provides a specialized agent framework for long-read single-cell transcriptomics.

## 2 Results

### 2.1 Overview of the scIsoAgent framework

We first constructed scIsoAgent as an interactive LLM agent for isoform-resolved long-read single-cell RNA-seq analysis. The framework was designed to support heterogeneous project entry points, ranging from raw long-read data to intermediate analysis results. This design reflects practical long-read single-cell projects, where users often need to continue from intermediate resources without being constrained by a fixed end-to-end pipeline.

scIsoAgent converts user-level analysis goals into executable and traceable workflow steps through four operational layers: request interpretation, skill selection, execution control, and interactive session management (Fig. 1). These layers implement three execution-oriented agent designs. Stage-aware hierarchical skill routing allows the agent to identify the analysis stage and dispatch requests to the appropriate workflow module. Script-based execution with user-confirmed commands keeps computational actions explicit and reproducible. Stateful tool reuse preserves interactive analysis contexts, such as loaded objects and active sessions, across related user requests. Together, these designs allow scIsoAgent to function as an agentic control layer connecting user intent, domain-specific skills, executable commands, and structured result interpretation.

**Figure 1:**
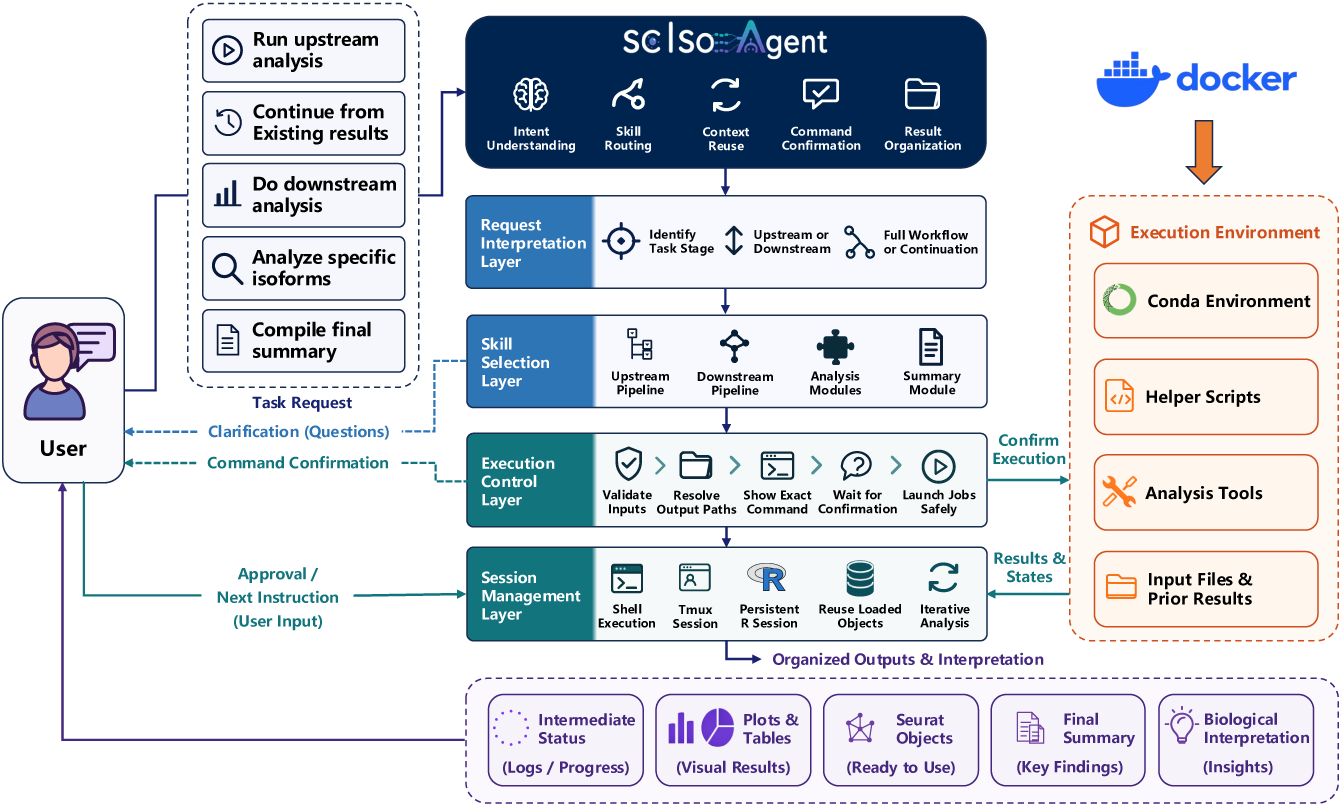
Overview of the scIsoAgent framework. scIsoAgent organizes long-read single-cell RNA-seq analysis as an interactive workflow that connects flexible user entry points with executable analysis and biological interpretation. User requests are interpreted and routed to stage-specific skills, including upstream processing, downstream single-cell analysis, isoform-level analysis, sequence-informed interpretation, and summary generation (top left). The execution control layer validates inputs, resolves output paths, and presents exact commands for user confirmation before launching jobs (center). The session management layer preserves computational context across iterative requests, enabling reuse of active sessions and loaded analysis objects (right). The framework interacts with containerized environments, helper scripts, analysis tools, input files, and prior results to generate traceable outputs and structured biological summaries (bottom).

This architecture allows users to move through long-read single-cell analysis as a guided, traceable interaction that can begin from different project stages and maintain continuity across iterative requests. In this way, scIsoAgent provides a flexible alternative to disconnected tool use or rigid static pipelines.

### 2.2 Workflow coverage for isoform-resolved long-read single-cell analysis

We next defined the workflow coverage of scIsoAgent across the major stages of isoform-resolved long-read single-cell analysis (Fig. 2). The workflow links platform-specific long-read processing to transcript-resolved single-cell analysis and sequence-informed interpretation within a unified agent-guided framework.

**Figure 2:**
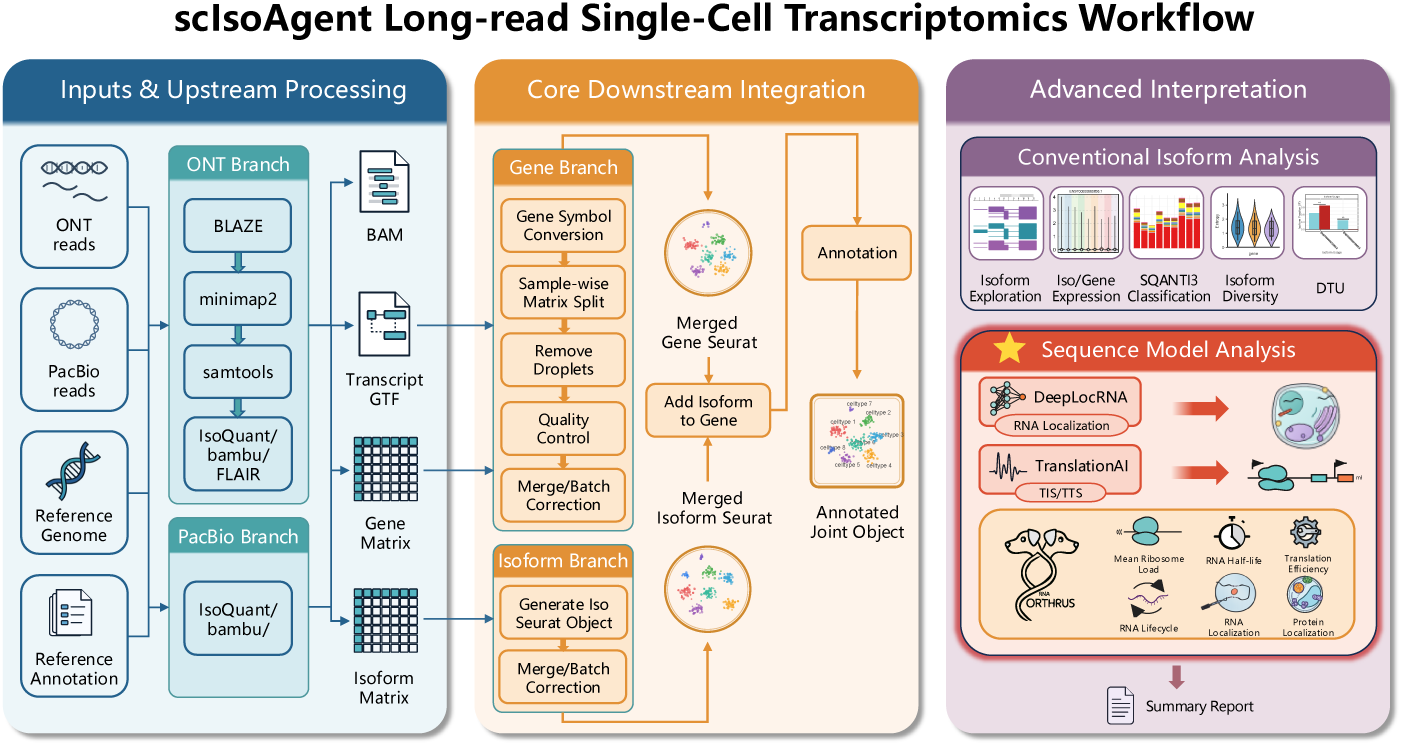
Workflow coverage of scIsoAgent for long-read single-cell RNA-seq analysis. The workflow is organized into three connected components. (1, left) Input and upstream processing support ONT and PacBio-based long-read single-cell data together with reference genome and annotation files. Platform-specific processing converts raw or intermediate inputs into standardized analysis resources, including aligned reads, transcript annotations, gene-level matrices, and isoform-level matrices. (2, middle) Core downstream integration uses gene-level and isoform-level matrices to construct single-cell analysis objects, perform quality control and integration, and link isoform-resolved information with cell-type annotations. This step generates annotated gene-level, isoform-level, and joint analysis objects for downstream exploration. (3, right) Advanced interpretation supports isoform exploration, expression visualization, transcript classification, isoform diversity analysis, differential transcript usage analysis, and sequence-informed interpretation. Sequence-level interpretation connects transcript sequences with RNA localization prediction, translation initiation and termination signal prediction, and transcript property prediction, followed by structured summary generation. DTU, differential transcript usage; TIS, translation initiation site; TTS, translation termination site.

scIsoAgent supports both ONT and PacBio-based workflows and can operate from raw sequencing data or processed long-read outputs. The workflow first harmonizes platform-dependent preprocessing and transcript quantification, then converts gene- and isoform-level outputs into inputs for downstream single-cell analysis. This organization allows transcript structures and isoform usage patterns to be interpreted together with cell identities and biological conditions, rather than as separate layers of analysis.

scIsoAgent further extends isoform-resolved analysis toward sequence-informed interpretation. In this component, scIsoAgent links transcript-level analysis results, including isoform expression, isoform classification, and differential transcript usage, with predictions from RNA sequence models. The interpretation module summarizes isoform-level evidence, structural annotations, and sequence-model predictions in a structured format, while explicitly distinguishing observed transcriptomic patterns from computationally inferred properties. This design supports biologically grounded interpretation of isoform-resolved findings and their potential sequence-level functional implications. The coordinated tool ecosystem underlying this workflow is summarized in Supplementary Fig. S2.

### 2.3 scIsoAgent improves planning quality and execution reliability

Because isoform-resolved long-read single-cell RNA-seq lacks a standardized end-to-end workflow, effective planning is essential for translating heterogeneous input resources and user-defined goals into executable analysis procedures. We therefore evaluated whether scIsoAgent could generate more appropriate analysis plans for this setting than general-purpose LLMs. We compared general-purpose LLM baselines with scIsoAgent variants built on different backbone models using the same long-read single-cell transcriptomic analysis prompt. The generated plans were assessed by LLM-based scoring and by specialists in computational biology.

In the automated planning evaluation, scIsoAgent variants achieved higher overall scores than general-purpose LLMs across three independent LLM-based evaluators, with scIsoAgent-GPT5.4 ranking highest (Fig. 3a). Dimension-level profiles showed that scIsoAgent generated more balanced plans across key criteria, supporting the benefit of domain-specific workflow structure for long-read single-cell analysis (Fig. 3b). The assessment by five specialists in computational biology further confirmed this trend. Compared with general-purpose LLMs, scIsoAgent variants received higher scores for domain-specific planning quality, including long-read single-cell specificity, executability, isoform resolution, and biological interpretability (Fig. 3c). At the group level, scIsoAgent variants significantly outperformed general-purpose LLMs in specialist-based assessment (Welch t-test, *p* = 1.66 × 10^−2^; Fig. 3d).

**Figure 3:**
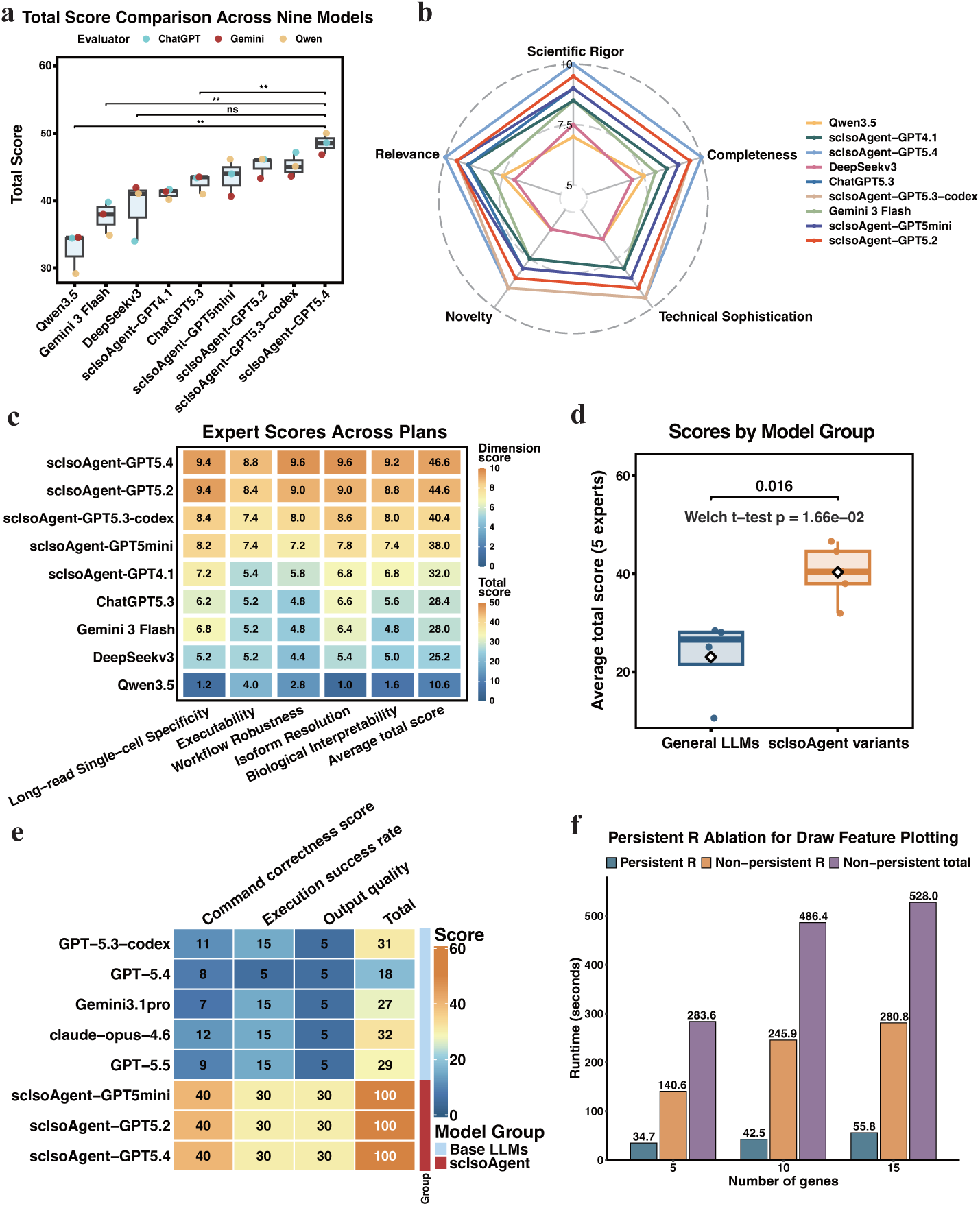
Evaluation of scIsoAgent planning and execution capabilities. **a,** Total scores of analysis plans generated by general-purpose LLMs and scIsoAgent variants across three LLM-based evaluators. Statistical comparisons were performed using Welch’s t-tests between scIsoAgent-GPT5.4 and each general-purpose LLM baseline shown by brackets. ns, not significant; **, *p <* 0.01. **b,** Radar plot showing dimension-level LLM-based planning scores across evaluated systems. Each plan was scored across five dimensions: scientific rigor, completeness, relevance, novelty, and technical sophistication. **c,** Heatmap of specialist-assigned mean scores across five domain-specific dimensions and the average total score. The five dimension scores are shown on a 0–10 scale, whereas the total score is shown on a separate 0–50 scale. **d,** Group-level comparison of specialist-assigned total scores between general-purpose LLMs and scIsoAgent variants. The center line indicates the median, boxes indicate the interquartile range, whiskers indicate the non-outlier range, and points indicate individual plans. Statistical significance was assessed using Welch’s t-test. **e,** IsoQuant-BAM execution evaluation based on command correctness, execution success rate, and output quality. **f,** Runtime comparison in the stateful tool-reuse evaluation for feature plotting with 5, 10, or 15 genes, comparing persistent R execution with non-persistent R execution and total non-persistent runtime.

We then tested whether improved planning translated into reliable execution using an IsoQuant-BAM task. This task required systems to construct a two-stage workflow from aligned ONT long-read single-cell BAM files, consisting of joint transcript discovery followed by per-sample single-cell quantification using the discovered transcript model. General-purpose LLMs often missed key workflow requirements, whereas all evaluated scIsoAgent variants achieved full scores for command correctness, execution success, and output quality (Fig. 3e). Finally, in an interactive feature-plotting task, stateful tool reuse reduced total runtime by maintaining a persistent R session and avoiding repeated object loading, with larger benefits as the number of requested genes increased (Fig. 3f).

Collectively, these evaluations indicate that scIsoAgent improves the continuity between analysis planning and computational execution, providing a domain-specific agent framework for reliable and interactive isoform-resolved long-read single-cell workflows.

### 2.4 Real-data application of scIsoAgent in a published long-read single-cell study

To evaluate whether scIsoAgent can support biologically meaningful analysis in a real study setting, we applied it to a published long-read single-cell study of the adult human heart and heart failure [29]. scIsoAgent guided a case-study reanalysis that linked cell-level organization with isoform-resolved transcript interpretation. The example prompts used for each analysis step and their corresponding reproduced output panels are summarized in Fig. 4a. This workflow progressed from cell integration and annotation to transcript classification, isoform distribution analysis, and differential transcript usage analysis.

**Figure 4:**
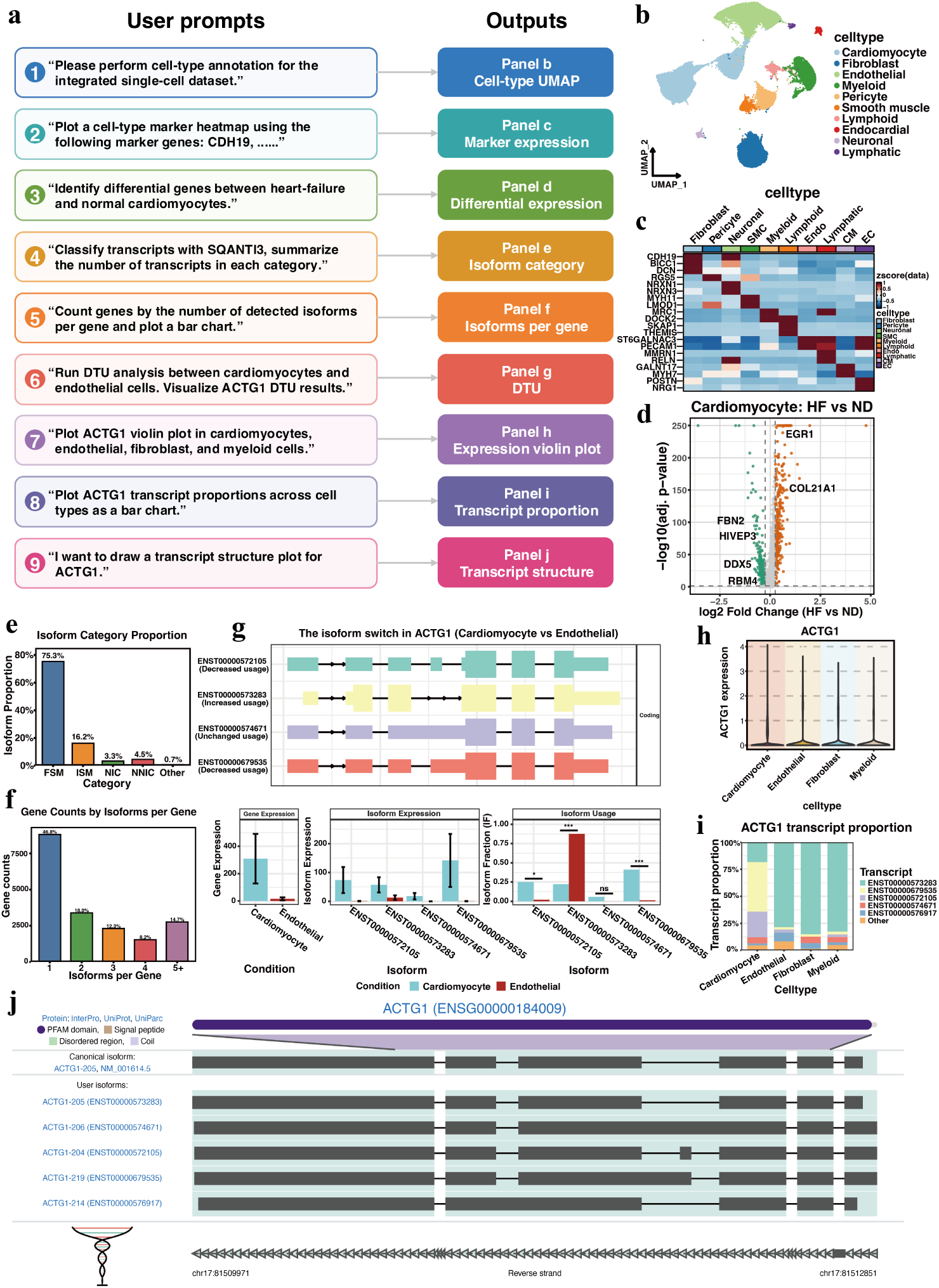
Reanalysis of a published long-read single-cell heart failure dataset using scIsoAgent. **a,** Prompt-to-analysis schematic showing how user prompts were parsed by scIsoAgent and converted into reproduced downstream analysis outputs. **b,** UMAP visualization of annotated major cardiac cell populations. **c,** The expression heatmap of marker genes across annotated cell types. **d,** Differential gene expression between heart failure and non-diseased cardiomyocytes. **e,** Isoform category proportions based on transcript classification. **f,** Distribution of genes by the number of detected isoforms per gene. **g,** Representative differential transcript usage event of *ACTG1* between cardiomyocytes and endothelial cells. Error bars indicate the standard error of the mean (SEM). **h,** *ACTG1* gene expression across selected major cell types. **i,** Transcript usage proportions of *ACTG1* across selected major cell types. **j,** Genome-track view showing the exon–intron structures and transcript annotations of representative *ACTG1* isoforms. EC, endothelial cells; CM, cardiomyocyte; HF, heart failure; ND, non-diseased; FSM, full splice match; ISM, incomplete splice match; NIC, novel in catalog; NNIC, novel not in catalog; DTU, differential transcript usage; ns, not significant; ***, *p <* 0.001.

scIsoAgent first reconstructed the cell-level organization of the dataset, which recovered the major cell populations reported in the original publication, including cardiomyocytes, fibroblasts, endothelial cells, pericyte, myeloid cells, neuronal cells, smooth muscle cells, lymphoid cells, endocardial cells, and lymphatic cells (Fig. 4b,c and Supplementary Fig. S3). Furthermore, scIsoAgent recovered reported differential gene expression patterns upon heart failure, including increase of *EGR1* and *COL21A1* and decrease of *HIVEP3* and *DDX5* in cardiomyocytes, increase of *PDE4D* and *FOS* and decrease of *FHL2* and *FRMD3* in endothelial cells, and increase of *POSTN* and *FGF14* and decrease of *ADGRB3* and *PLA2G5* in fibroblasts (Fig. 4d and Supplementary Fig. S4a,b). These results show that scIsoAgent can recover both cellular organization and disease-associated expression patterns from long-read single-cell resources.

At the transcript level, scIsoAgent recovered the isoform-resolved landscape reported in the original study. Detected transcripts spanned known and novel structural classes, including full-splice match (FSM), incomplete-splice match (ISM), novel in catalog (NIC), and novel not in catalog (NNIC) isoforms (Fig. 4e). scIsoAgent also reproduced the expected distribution of detected isoforms per gene, in which most genes were represented by a single isoform and progressively fewer genes showed multiple isoforms (Fig. 4f).

scIsoAgent further recovered representative differential transcript usage events associated with cellular identity and disease status. For *ACTG1*, the reanalysis reproduced cell-type-associated isoform usage differences between cardiomyocytes and other major cell types, with transcript structure views linking these differences to specific isoforms (Fig. 4g–j and Supplementary Fig. S5a,b). Additional examples included cardiomyocyte-associated DTU of *TNNI3* and heart-failure-associated DTU of *FRY* in cardiomyocytes (Supplementary Fig. S5c-k). Together, these results show that scIsoAgent can recover both global isoform features and representative transcript-usage changes from a real long-read single-cell study.

Overall, this case study demonstrates that scIsoAgent can coordinate cell-level, gene-level, isoform-level, and DTU-focused analyses from existing long-read single-cell resources, supporting its practical use in real biological reanalysis settings.

### 2.5 Sequence-informed interpretation of isoform function

Long-read single-cell RNA-seq links transcript sequences with cell-type-specific isoform expression and usage patterns, creating an opportunity to interpret isoform variation beyond detection and quantification. However, isoform-level observations do not by themselves reveal whether alternative transcripts differ in sequence-associated properties such as stability, localization, or translation potential. scIsoAgent addresses this interpretive gap by connecting observed isoform evidence with predictions derived from RNA sequence models. This design enables sequence-informed interpretation of isoform variation while preserving a clear distinction between measured transcriptomic evidence and model-inferred functional properties.

We next used scIsoAgent to extend isoform-resolved single-cell analysis toward sequence-informed interpretation. We selected *TNNI3*, a cardiomyocyte-associated DTU event identified in the preceding analysis, as a representative case. scIsoAgent extracted the corresponding isoform sequences and evaluated whether the major transcript TNNI3-201 and the intron-retention-associated transcript TNNI3-206 differed in predicted transcript behavior. This analysis focused on three complementary properties: translational and stability-related potential predicted by Orthrus [30], RNA localization predicted by DeepLocRNA [31], and translation-related sequence signals predicted by TranslationAI [32].

Sequence-model outputs suggested that TNNI3-201 (ENST00000344887) and TNNI3-206 (ENST00000587176) differ in predicted transcript productivity and localization tendency. TNNI3-201 showed stronger predicted translation efficiency, mean ribosome load, and RNA stability-related signals than TNNI3-206, suggesting that the major transcript may represent a more productive and stable isoform state (Fig. 5a). TNNI3-201 also showed an Exosome-associated localization signal that was not observed for TNNI3-206 (Fig. 5b). Both transcripts retained ORF-compatible initiation and termination signals, indicating that TNNI3-206 is not simply a non-translatable transcript despite its weaker predicted translational and stability-related profile (Fig. 5c).

**Figure 5:**
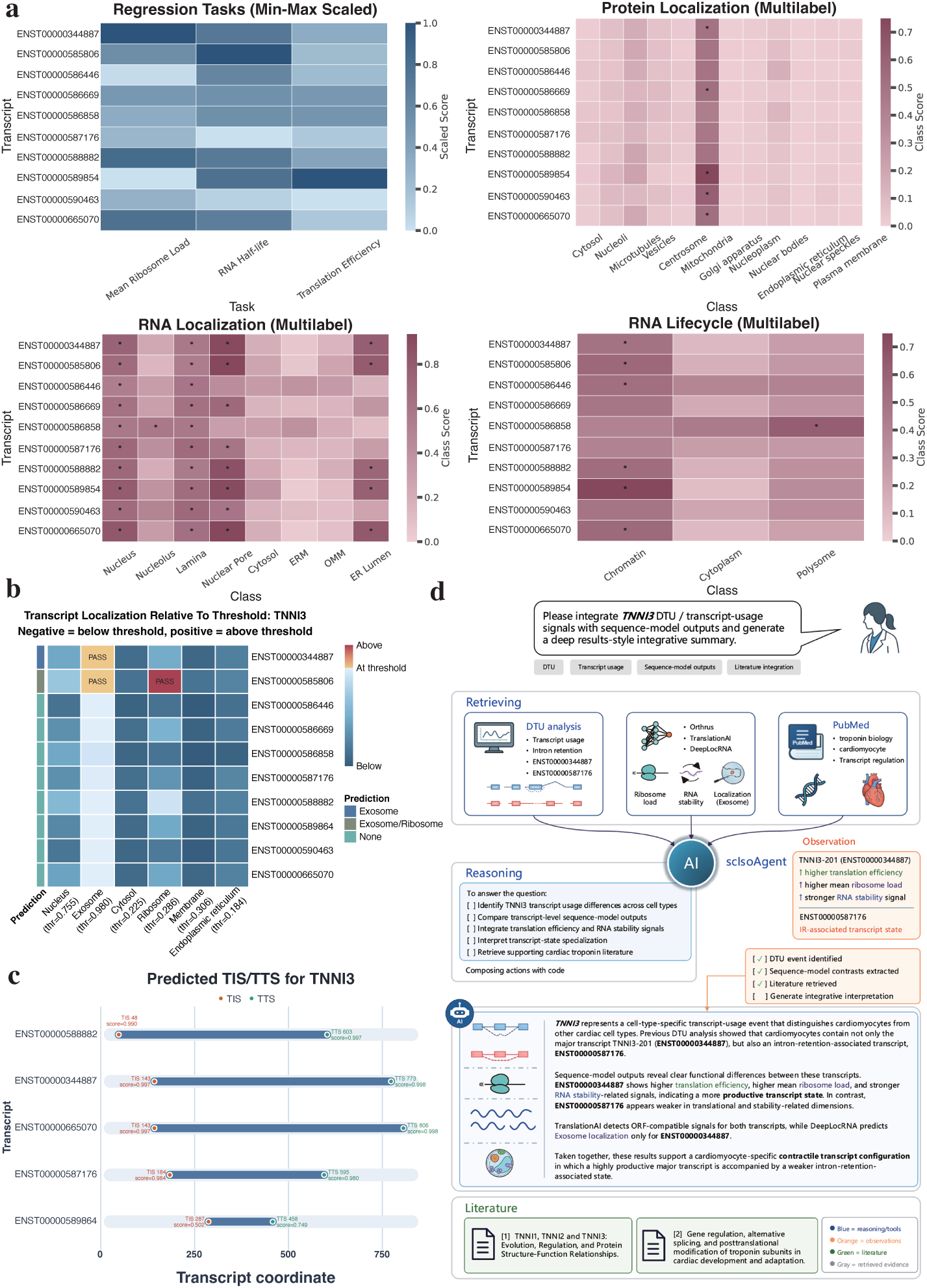
Sequence-informed interpretation of a *TNNI3* transcript-usage event using scIsoAgent. **a,** Orthrus-based predictions for *TNNI3* transcripts, including regression outputs for mean ribosome load, RNA half-life, and translation efficiency, together with multilabel predictions related to protein localization, RNA localization, and RNA lifecycle-associated classes. **b,** DeepLocRNA-based transcript localization predictions for *TNNI3* transcripts. Values are shown relative to model-specific localization thresholds, with negative values below the threshold and positive values above the threshold. **c,** TranslationAI-based prediction of translation initiation sites and translation termination sites across selected *TNNI3* transcripts. TIS, translation initiation site; TTS, translation termination site. **d,** scIsoAgent-generated interpretation summary for the representative *TNNI3* DTU event. The panel shows the agentic interpretation process, including evidence retrieval, reasoning, observation extraction, summary generation, and literature contextualization. In the retrieving stage, scIsoAgent collects DTU evidence, transcript-usage results, sequence-model outputs, and relevant literature context. The reasoning and observation modules organize these inputs into explicit intermediate conclusions, including the contrast between TNNI3-201 and TNNI3-206. The generated summary integrates observed transcript-usage evidence with model-inferred transcript properties to produce a sequence-informed biological interpretation, while the literature module provides supporting context for cardiac troponin biology. DTU, differential transcript usage.

scIsoAgent then integrated the DTU evidence, sequence-derived predictions, and literature-oriented context into a structured interpretation. The generated summary described a cardiomyocyte-associated *TNNI3* transcript-usage pattern in which the major transcript TNNI3-201 appeared to represent a more productive isoform state, whereas TNNI3-206 represented an intron-retention-associated isoform with weaker predicted translational and stability-related properties but retained ORF-compatible signals. This example illustrates how scIsoAgent can transform an isoform-level single-cell observation into a sequence-informed functional hypothesis while maintaining a clear distinction between measured transcript usage and model-inferred transcript properties (Fig. 5d).

Together, this *TNNI3* case study shows that scIsoAgent can connect observed isoform usage with model-inferred transcript properties, extending long-read single-cell analysis toward sequence-informed, hypothesis-generating interpretation.

## 3 Discussion

In this study, we developed scIsoAgent, an LLM-powered agent for isoform-resolved long-read single-cell RNA-seq analysis. Long-read single-cell transcriptomics provides transcript structures, isoform diversity, transcript usage, and sequence-level information that are difficult to recover from conventional short-read single-cell data [33]. At the same time, these information layers create a coordination challenge: transcript-level evidence must be connected with cellular context, biological condition, and sequence-informed interpretation across heterogeneous computational resources. scIsoAgent addresses this challenge by organizing long-read single-cell analysis as an interactive and executable workflow that connects heterogeneous inputs with isoform-level analysis and biological interpretation.

A key insight from scIsoAgent is that long-read single-cell analysis is not only a reasoning problem, but also a workflow-coordination problem. This perspective distinguishes scIsoAgent from broader biomedical [26, 28] or autonomous research agents [34, 35], which often emphasize literature exploration, hypothesis generation, or general omics support. In contrast, scIsoAgent focuses on a narrower but technically demanding setting in which useful biological interpretation depends on coordinating platform-specific processing, transcript-level evidence, single-cell context, and executable downstream analysis. By combining stage-aware workflow control, explicit script-based execution, and reusable computational context, scIsoAgent provides an agentic layer that links user goals to domain-specific analysis steps while preserving traceability and reproducibility.

Our evaluations support this design principle without relying only on benchmark prompts. scIsoAgent generated more domain-aligned plans than general-purpose LLM baselines, translated workflow decisions more reliably into executable analysis steps, and improved iterative analysis through stateful tool reuse. In a real-data reanalysis, it further coordinated cell-level and isoform-level analyses from published long-read single-cell resources. These results suggest that scientific agents are particularly useful for specialized omics settings when they combine domain knowledge, executable workflows, and persistent analysis context rather than simply producing free-form analytical suggestions. A further contribution of scIsoAgent is its ability to connect isoform-resolved analysis with sequence-informed interpretation. Recent nucleotide and RNA sequence models have expanded the possibility of inferring regulatory, localization, translation-related, and other functional properties from sequence [36, 37, 38, 39, 40, 30, 31, 32]. However, these models are often applied as standalone prediction tools, disconnected from cell-type-specific or condition-specific isoform evidence. scIsoAgent provides a framework for linking transcript-level observations with sequence-derived predictions and structured interpretation. In the *TNNI3* example, it integrated DTU evidence with Orthrus, DeepLocRNA, and TranslationAI outputs to generate a hypothesis about isoform-level differences in transcript behavior. Such predictions require experimental validation, but they offer a principled way to prioritize isoforms for downstream functional studies.

This study also has several limitations. Our execution benchmarks cover representative tasks rather than the full diversity of long-read single-cell workflows, and future evaluations should include broader input types, platforms, and biological use cases. Both automated and specialist-based planning assessments may be influenced by the selected prompts, scoring rubrics, and evaluator backgrounds. Finally, sequence-derived predictions should be interpreted as hypothesis-generating evidence rather than direct functional validation.

In summary, scIsoAgent provides an autonomous agent framework for isoform-resolved long-read single-cell transcriptomics. It links adaptive workflow planning, traceable execution, and sequence-informed interpretation within a coherent analysis process. More broadly, this study highlights the value of scientific agents as reproducible workflow coordinators for specialized biomedical analyses.

## 4 Methods

### 4.1 Implementation of scIsoAgent

The overall procedure of scIsoAgent is summarized in Algorithm 1. scIsoAgent was implemented as an execution-aware agent for isoform-resolved long-read single-cell RNA-seq analysis. Given a user request, project context, available input resources, and an active session state, scIsoAgent first interprets the user goal and identifies the corresponding analysis stage. It then routes the request to a stage-specific skill module, validates the required inputs, selects the appropriate software environment, and generates an executable command or script call. Before execution, the command is displayed to the user and launched only after confirmation. After execution, scIsoAgent collects outputs, updates the reusable session state, and generates structured result summaries or biological interpretations. This procedure allows scIsoAgent to support both end-to-end workflows and continuation from intermediate resources while preserving traceability and reproducibility across interactive analysis steps.

**Algorithm 1**. scIsoAgent execution procedure for long-read single-cell transcriptome analysis

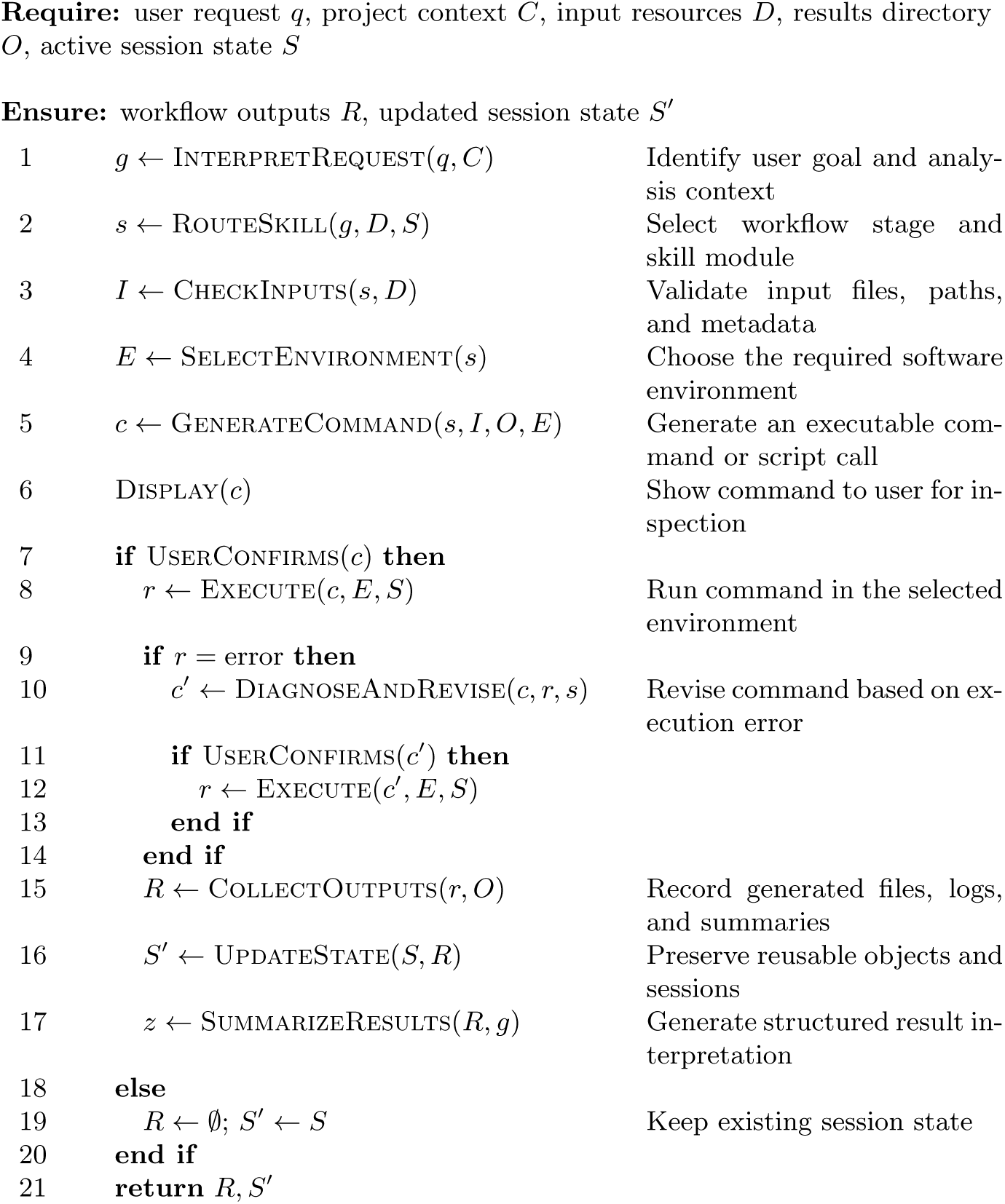

The implementation follows this procedure through a portable workflow bundle organized into two sibling directories, skills/ and scripts/. The skills/ layer contains agent-facing modules that encode stage-aware routing, required inputs, user interaction rules, execution policies, and output organization. A top-level routing skill assigns user requests to the appropriate workflow stage, including upstream processing, downstream single-cell analysis, isoform-level analysis, sequence-informed interpretation, or summary generation. The scripts/ layer contains executable R and Python helper scripts for standardized computational steps. This separation allows scIsoAgent to keep workflow logic and execution policy in the skills layer while delegating computation to inspectable helper scripts.

scIsoAgent was designed to make computational actions explicit and reusable. For substantial computational steps, the agent generates the exact command, displays it for user inspection, and proceeds only after confirmation. Before execution, it validates input paths, resolves output locations, checks required resources, and avoids silently overwriting existing outputs. The runtime environment is Docker-based, with workflow stages assigned to predefined conda environments and outputs organized under a user-confirmed results directory. To support stateful tool reuse, long-running command-line tasks can be launched in tmux sessions, and interactive R analyses can use persistent R sessions so that loaded Seurat objects and intermediate analysis states are retained across related requests.

### 4.2 Workflow organization and supported entry points

scIsoAgent organizes isoform-resolved long-read single-cell RNA-seq analysis into modular workflow stages rather than a fixed linear pipeline. These stages cover platform-specific upstream processing, transcript quantification, downstream single-cell integration, cell-type annotation, isoform-level analysis, sequence-informed prediction, and summary generation. Each stage can be executed independently or linked to other stages according to the available input resources and the user’s analysis goal.

The workflow supports flexible entry points that reflect common long-read single-cell analysis scenarios. Users can begin from raw ONT or PacBio reads, aligned BAM files, transcript annotations, gene- or isoform-level expression matrices, transcript FASTA files, merged single-cell objects, or selected genes and isoforms for targeted interpretation. At the start of each request, scIsoAgent checks whether the required inputs are available and asks for missing resources before generating executable commands. Validated paths, output directories, and intermediate results are retained as reusable context, allowing users to continue from intermediate or published resources without restarting a complete end-to-end analysis.

### 4.3 Upstream read processing and transcript quantification

The upstream module converts long-read single-cell RNA-seq inputs into transcript-level resources for downstream analysis. Required inputs include long-read sequencing files or preprocessed alignments, reference genome and annotation files, sample identifiers, an output directory, and the sequencing platform. scIsoAgent validates these inputs, determines whether the workflow should follow an ONT- or PacBio-compatible route, and generates the corresponding executable commands.

For ONT data, scIsoAgent supports barcode and UMI recovery with BLAZE [41], followed by genome alignment using minimap2 [42]. When barcode or UMI tags are available, they are preserved for downstream single-cell quantification. Alignment files are then processed with samtools for sorting, indexing, and validation [43]. For PacBio CCS data, scIsoAgent starts from platform-compatible long-read inputs or alignments and does not use BLAZE-based barcode and UMI recovery as the default preprocessing route.

Transcript discovery and quantification are performed using user-selected long-read transcriptomic tools, including IsoQuant [44], bambu [45], or FLAIR [46]. scIsoAgent selects or requests the quantification route according to the platform, available input files, and analysis goal. IsoQuant and bambu workflows generate transcript models and expression matrices from compatible long-read inputs. For FLAIR-based workflows, scIsoAgent supports transcriptome construction, quantification, and generation of single-cell gene- and transcript-level matrices through project helper scripts.

The main outputs of the upstream module include processed alignment files, transcript annotations, gene- and isoform-level expression matrices, and execution logs or summary statistics. These outputs are organized under the user-confirmed results directory and passed forward as reusable resources for downstream single-cell integration and isoform-level analysis.

### 4.4 Core downstream integration of gene- and isoform-level expression

The downstream integration module converts quantified long-read outputs into single-cell objects that preserve both gene-level and isoform-level information. Supported inputs include expression matrices, transcript annotations, reference annotations, or existing Seurat objects. scIsoAgent checks the available resources, identifies the appropriate entry point, and generates scripts for object construction and integration.

scIsoAgent first builds a transcript-to-gene mapping table to link isoform identifiers with their corresponding genes. Depending on the input type, this mapping is derived from quantification outputs, transcript annotations, or existing isoform matrices. The resulting table provides a shared reference for isoform analysis, gene–isoform integration, and sequence-based interpretation.

For gene-level processing, scIsoAgent constructs sample-level Seurat objects and applies standard single-cell quality control. Empty droplets are removed using DropletUtils when required [47], and doublets are detected with DoubletFinder when applicable [48]. User-confirmed parameters control filtering, dimensionality reduction, clustering, and doublet detection. After quality control, scIsoAgent merges the sample-level objects and processes them with Seurat for normalization, dimensionality reduction, clustering, and UMAP visualization [49]. SCP is used for selected visualization and downstream utilities [50], and Harmony-based batch correction can be applied after user confirmation [51].

Isoform-level matrices are processed in parallel and aligned to the retained cell barcodes in the matched gene-level object. scIsoAgent then adds isoform counts as a separate assay in the merged Seurat object, allowing gene-level cell annotations and isoform-level measurements to be analyzed in the same framework. Cell-type annotation is performed using marker-based strategies or optional ScType-assisted annotation [52]. The module produces reusable integrated Seurat objects and associated quality-control, annotation, and visualization outputs for downstream isoform analysis. The workflow organization was informed by FLAMESv2, a full-length isoform analysis framework for single-cell and spatial RNA-seq data [53].

### 4.5 Isoform-level characterization and transcript usage analysis

Isoform-level characterization is performed after transcript quantification and gene–isoform integration. Supported inputs include isoform expression data, transcript annotations, SQANTI3 classification results, transcript-to-gene mappings, or Seurat objects containing an isoform assay. Before execution, scIsoAgent validates the required resources and confirms the metadata fields, grouping variables, filtering parameters, and output locations.

For isoform exploration, scIsoAgent aggregates transcript-level expression across user-specified groups, such as cell type, sample, condition, or cluster. Transcript identifiers are linked to gene identifiers and gene symbols so that isoform usage can be summarized at both transcript and gene levels. This allows users to inspect selected genes, compare isoform expression across groups, and identify genes with complex transcript usage patterns.

Isoform complexity is summarized using the number of detected transcripts per gene and entropy-based measures of within-gene transcript usage diversity. These analyses produce gene-level summaries and visualization outputs that highlight genes or cell populations with high isoform diversity.

For structural characterization, scIsoAgent supports SQANTI3-based transcript classification using a reference genome and annotation [54]. SQANTI3 categories, including full-splice match, incomplete-splice match, novel in catalog, and novel not in catalog transcripts, are linked back to expression matrices through transcript identifiers. This step allows structural transcript classes to be analyzed together with cell-type- or condition-specific expression patterns.

Differential transcript usage analysis is performed from isoform-level counts. When required, scIsoAgent aggregates transcript counts into pseudobulk matrices according to the selected biological grouping and applies IsoformSwitch-AnalyzeR to evaluate transcript switching or differential usage patterns [55]. Selected events can be visualized using isoform usage plots, switch-style summaries, sashimi plots generated by ggsashimi [56], or transcript structure views generated by IsoVis [57]. The resulting tables, plots, and analysis objects are organized as reusable outputs for downstream sequence-informed interpretation and reporting.

### 4.6 Sequence-based isoform functional prediction and interpretation

scIsoAgent supports sequence-based isoform interpretation using full-length transcript sequences. The required inputs are transcript FASTA files and a transcript-to-gene mapping table. When transcript FASTA files are not available, scIsoAgent generates them from transcript annotations and a reference genome using gffread [58]. For gene-specific analysis, the agent extracts all transcripts assigned to the selected gene and writes them to a gene-specific FASTA file for downstream prediction.

scIsoAgent applies three RNA sequence-model modules to the extracted isoform sequences. Orthrus is used to predict transcript-level properties related to translation, RNA stability, localization, and lifecycle features [30]. DeepLocRNA is used to predict RNA subcellular localization profiles [31]. TranslationAI is used to predict translation initiation and termination-related signals [32]. The resulting predictions are organized into isoform-level tables and visualizations, including heatmaps and transcript-coordinate plots when applicable. All model outputs are treated as computational predictions rather than direct experimental measurements.

The sequence-based prediction module can be linked to isoform-level evidence generated by earlier scIsoAgent workflows, including differential transcript usage analysis, isoform exploration, and isoform diversity profiling. For interpretation, scIsoAgent combines observed isoform evidence, transcript annotations, sequence-derived predictions, and generated outputs into a structured report. The report explicitly distinguishes measured expression or usage patterns from model-inferred transcript properties and is used to generate hypothesis-oriented functional interpretations of selected isoforms.

### 4.7 PubMed-based literature survey

A PubMed-based literature survey was performed to summarize the development of long-read single-cell transcriptomics over time and to characterize its associated information and method landscape. Annual publication counts were obtained using the NCBI PubMed ESearch utility. Searches were restricted to title and abstract fields, stratified by publication year, and summarized as non-cumulative yearly hit counts from 2009 to 2025. Review articles were excluded using the keyword term NOT review.

For publication trends, short-read single-cell RNA-seq queries combined terms for single-cell or single-nucleus RNA sequencing. Long-read single-cell RNA-seq queries further included long-read platform and transcriptomic terms. To compare growth trajectories, observed long-read single-cell publication years were shifted seven years earlier on the x-axis relative to their original publication years. The dashed curve was used as a heuristic visual projection based on the earlier growth pattern of short-read single-cell RNA-seq, and was not treated as observed PubMed counts.

To summarize information content, the long-read single-cell base query was combined with keyword groups representing isoform structure, splicing, transcript boundaries, allele- or haplotype-related information, fusion transcripts, and RNA-level signals. To summarize the method ecosystem, the same base query was combined with keyword groups representing experimental protocols, computational pipelines, benchmarking, quality control, and short-read integration. Category counts were not mutually exclusive. Exact ESearch URLs, query terms, keyword groups, and annual counts were retained as source tables.

### 4.8 Planning evaluation

We evaluated analysis-plan generation using the same prompt for all systems. The evaluated systems included four general-purpose LLMs, Qwen3.5, Gemini 3 Flash, DeepSeekv3, and ChatGPT5.3, and five scIsoAgent variants, scIsoAgent-GPT4.1, scIsoAgent-GPT5mini, scIsoAgent-GPT5.2, scIsoAgent-GPT5.3-codex, and scIsoAgent-GPT5.4. The scIsoAgent variants used the same domain-specific skill framework but differed in the underlying language model or agent configuration. General-purpose LLMs were evaluated without the scIsoAgent framework.

Generated plans were evaluated under two complementary scoring settings. In both settings, evaluators were blinded to the system labels associated with each generated plan. In the automated setting, three LLM-based evaluators, ChatGPT5.5, Gemini 3 Flash, and Qwen3.5, independently scored each plan using a five-dimension rubric covering scientific quality, completeness, relevance, novelty, and technical sophistication. In the specialist setting, five computational biology specialists with 3–5 years of experience in single-cell or computational transcriptomic analysis scored each plan using a domain-specific rubric covering long-read single-cell relevance, executability, workflow robustness, isoform resolution, and biological interpretability. In both settings, each dimension was scored on a 10-point scale, and total scores were calculated as the sum of the five dimensions, with a maximum of 50. Specialist scores were averaged across evaluators for each system. Group-level differences between general-purpose LLMs and scIsoAgent variants were tested using Welch’s t-test.

### 4.9 Execution evaluation

We performed two execution-oriented evaluations to assess command reliability and stateful tool reuse. In the IsoQuant-BAM task, we compared five general-purpose LLMs, GPT-5.3-codex, GPT-5.4, Gemini 3.1 Pro, Claude Opus 4.6, and GPT-5.5, with three scIsoAgent variants, scIsoAgent-GPT5mini, scIsoAgent-GPT5.2, and scIsoAgent-GPT5.4. Each system was asked to generate and execute the expected IsoQuant workflow from two aligned ONT long-read single-cell BAM files. The expected solution required a two-stage strategy: joint transcript discovery using both BAM files, followed by per-sample single-cell quantification using the discovered transcript model with model construction disabled.

Outputs were evaluated using a 100-point rubric covering command correctness, execution success, and output quality. Command correctness contributed 40 points and assessed whether the generated commands followed the required two-stage IsoQuant workflow with appropriate parameters. Execution success contributed 30 points and was scored according to the number of correction rounds required to complete the task. Output quality contributed 30 points and assessed whether the expected files were generated and matched the intended workflow.

The stateful tool-reuse evaluation used an interactive feature-plotting task based on a Seurat object. We compared a persistent setting, in which the object was loaded once and reused in an active R session, with a non-persistent setting, in which the same RDS file was reloaded for each plotting request. Runtime was measured for plotting 5, 10, and 15 genes.

### 4.10 Reanalysis and sequence-informed extension of a published long-read single-cell RNA-seq dataset

We evaluated scIsoAgent on a published long-read single-cell RNA-seq dataset from the Gene Expression Omnibus under accession number GSE288222 [29]. The user provided scIsoAgent with the released quantified resources from the original study, rather than raw sequencing reads. These resources included a transcript-quantification-derived GTF file, a gene expression matrix, and a transcript/isoform expression matrix. The user also specified the biological comparison of interest as heart failure versus normal samples.

Starting from these inputs, scIsoAgent identified the available resource types and selected the corresponding reanalysis workflow. It generated and executed scripts to construct gene-level and isoform-level analysis objects from the provided matrices. The gene expression matrix was used to support dimensionality reduction, clustering, UMAP visualization, cell-type annotation, marker visualization, and cell-type-specific differential gene expression analysis. The transcript/isoform expression matrix and transcript-derived GTF file were used for isoform-level analysis, transcript-to-gene mapping, transcript structural annotation, and differential transcript usage analysis. For disease-associated gene-level comparisons, scIsoAgent performed differential expression analysis between heart failure and normal samples within major cardiac cell types, including cardiomyocytes, endothelial cells, and fibroblasts.

For isoform-level reanalysis, scIsoAgent generated scripts to assign transcript structural categories using SQANTI3 and to summarize isoform features across annotated cell types. These analyses included isoform category proportions, the distribution of genes by the number of detected isoforms per gene, and representative cell-type- or condition-associated differential transcript usage events. The resulting outputs were compared with findings reported in the original study to assess whether scIsoAgent could recover major cell-level and isoform-level patterns from released quantified resources.

For sequence-informed extension, the user selected a representative DTU-associated gene for transcript-level functional interpretation. scIsoAgent extracted the relevant transcript sequences from transcript FASTA resources or generated them from transcript annotation and reference genome files when needed. It then applied RNA sequence models to evaluate transcript-level properties, RNA localization, and translation initiation or termination-related signals. Finally, scIsoAgent organized the DTU evidence, sequence-derived predictions, and transcript-level context into an interpretation-oriented summary, while maintaining a distinction between observed transcriptomic evidence and model-inferred properties.

## 5 Code availability

The codebase for scIsoAgent is publicly available at https://github.com/zczali4403/scIsoAgent. The repository contains the agent-facing skill modules, executable helper scripts, documentation, and example prompts for running the scIsoAgent workflow.

## Acknowledgements

This work was supported by the National Key Research and Development Program of China (No. 2024YFF1207500 to Y.X.) and the National Natural Science Foundation of China (No. 32470703 and 23DAA01060 to Y.X.). We thank the five anonymous specialists in computational biology for their assistance with the domain-specific planning assessment.

## Supplementary Figures

**Figure S1:**
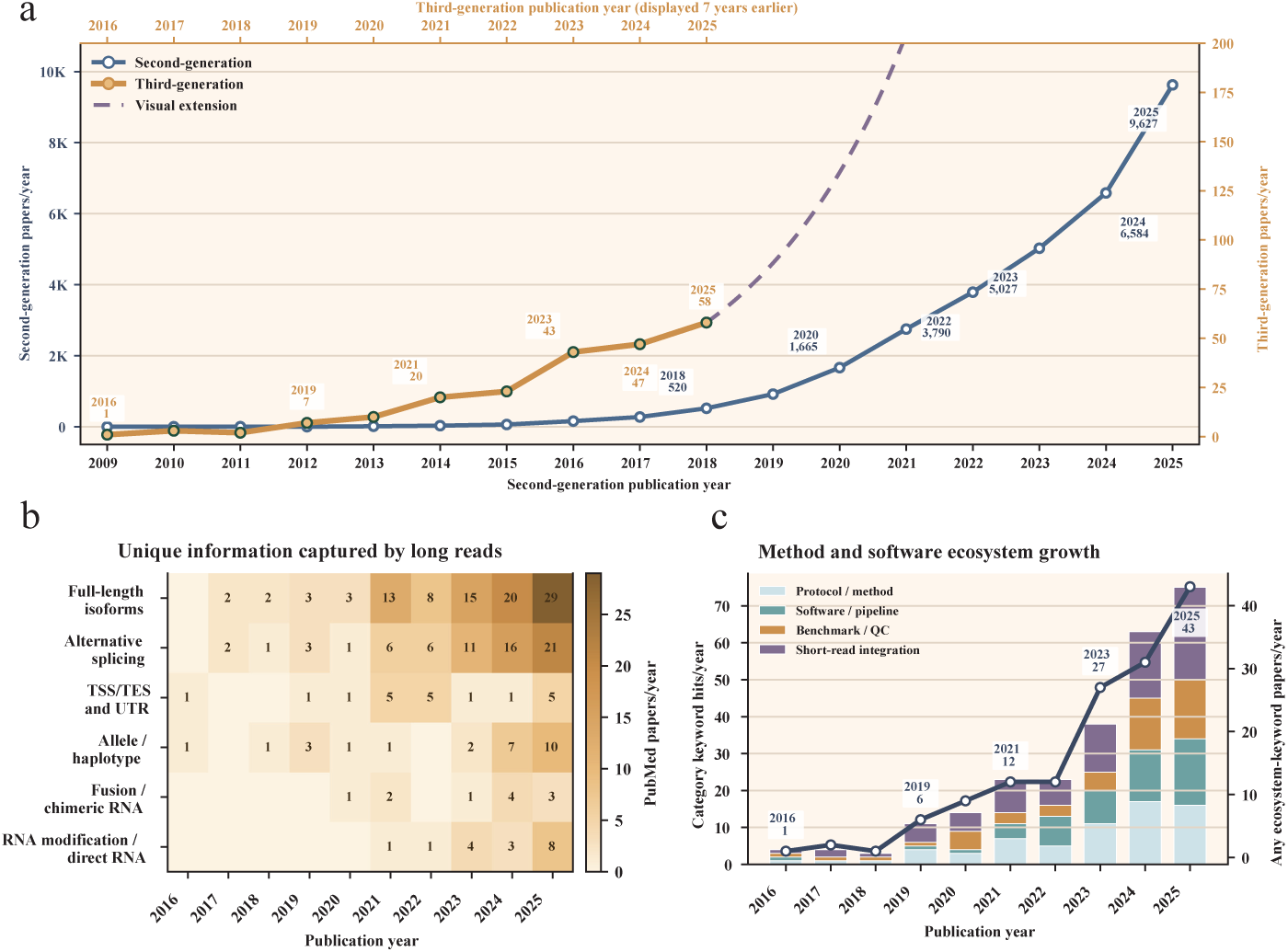
PubMed-based literature survey of long-read single-cell transcriptomics. **a,** Annual publication trends for second-generation and third-generation single-cell RNA-seq studies. Third-generation studies are shown on a separate right y-axis. The dashed curve indicates the projected future growth of third-generation single-cell studies, estimated from the earlier growth trajectory of second-generation single-cell RNA-seq studies after a 7-year temporal shift. **b,** Yearly PubMed keyword-hit trends for information types enabled or improved by long-read sequencing. **c,** Yearly PubMed keyword-hit trends for method and software ecosystem categories. Keyword-hit counts are survey-level summaries and are not mutually exclusive publication categories. TSS, transcription start site; TES, transcription end site; UTR, untranslated region; QC, quality control.

**Figure S2:**
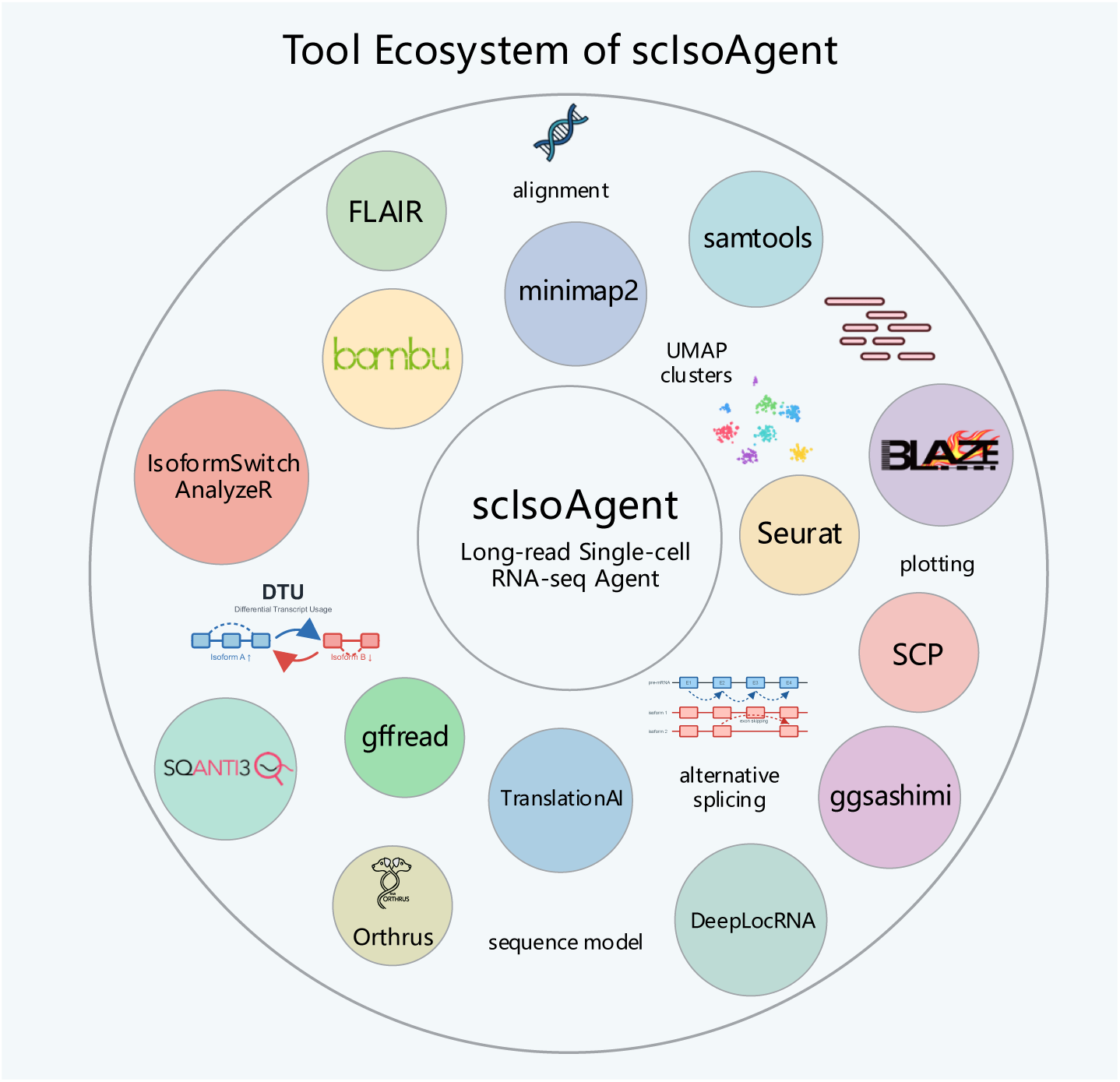
Tool ecosystem coordinated by scIsoAgent for long-read single-cell RNA-seq analysis. The schematic shows representative tools and analysis modules organized around scIsoAgent, spanning long-read preprocessing, single-cell integration and visualization, isoform-level analysis, differential transcript usage interpretation, and sequence-informed prediction.

**Figure S3:**
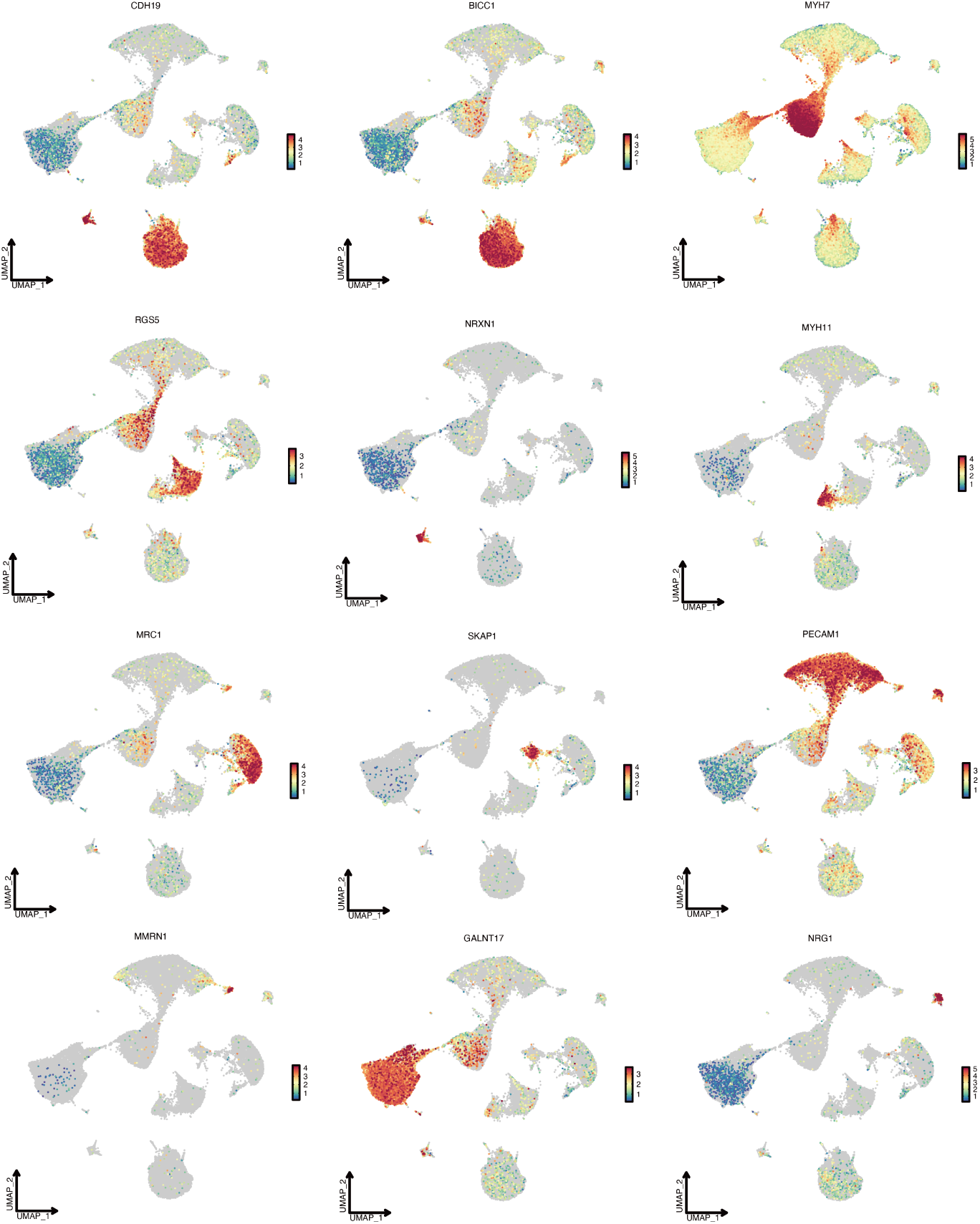
The expression of marker genes in the single-cell dataset (GSE288222). UMAP feature plots showing marker genes for major cell populations, including lymphoid cells, fibroblasts, cardiomyocytes, neuronal cells, smooth muscle cells, endothelial cells, myeloid cells, endocardial cells, and lymphatic cells.

**Figure S4:**
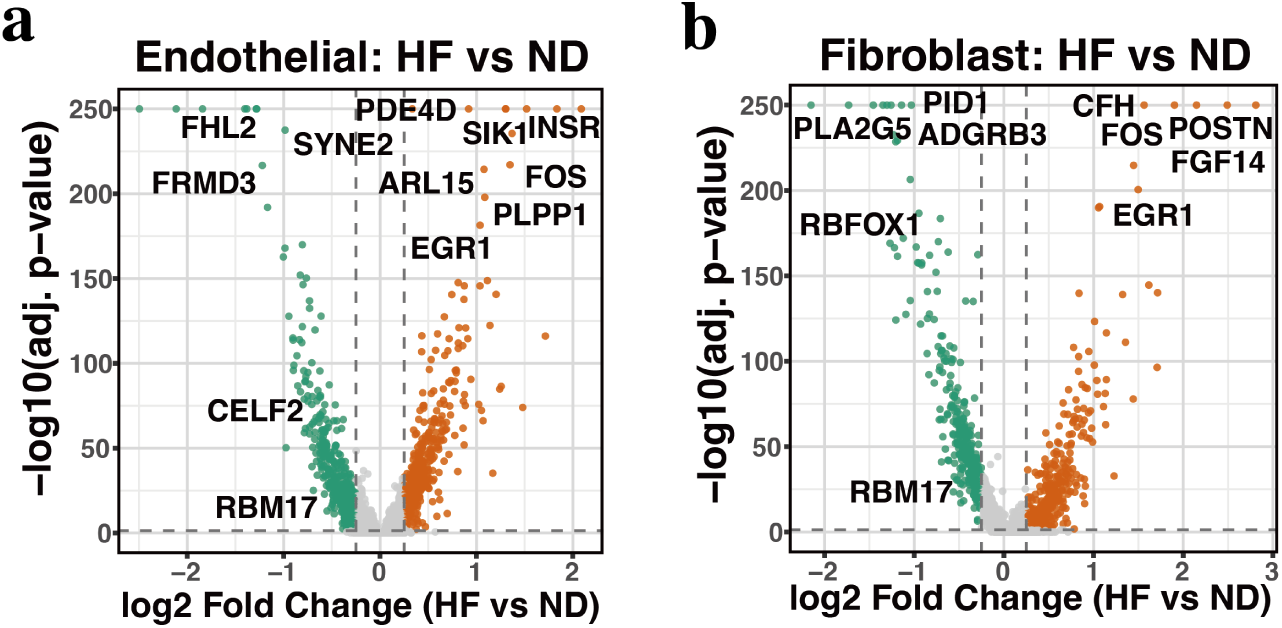
Differential gene expression between heart failure and non-diseased cell types. **a,** Volcano plot showing differentially expressed genes between heart failure and non-diseased endothelial cells. **b,** Volcano plot showing differentially expressed genes between heart failure and non-diseased fibroblasts. The y-axis shows − log_10_ adjusted *p* values calculated using the Mann–Whitney *U* test. HF, heart failure; ND, non-diseased.

**Figure S5:**
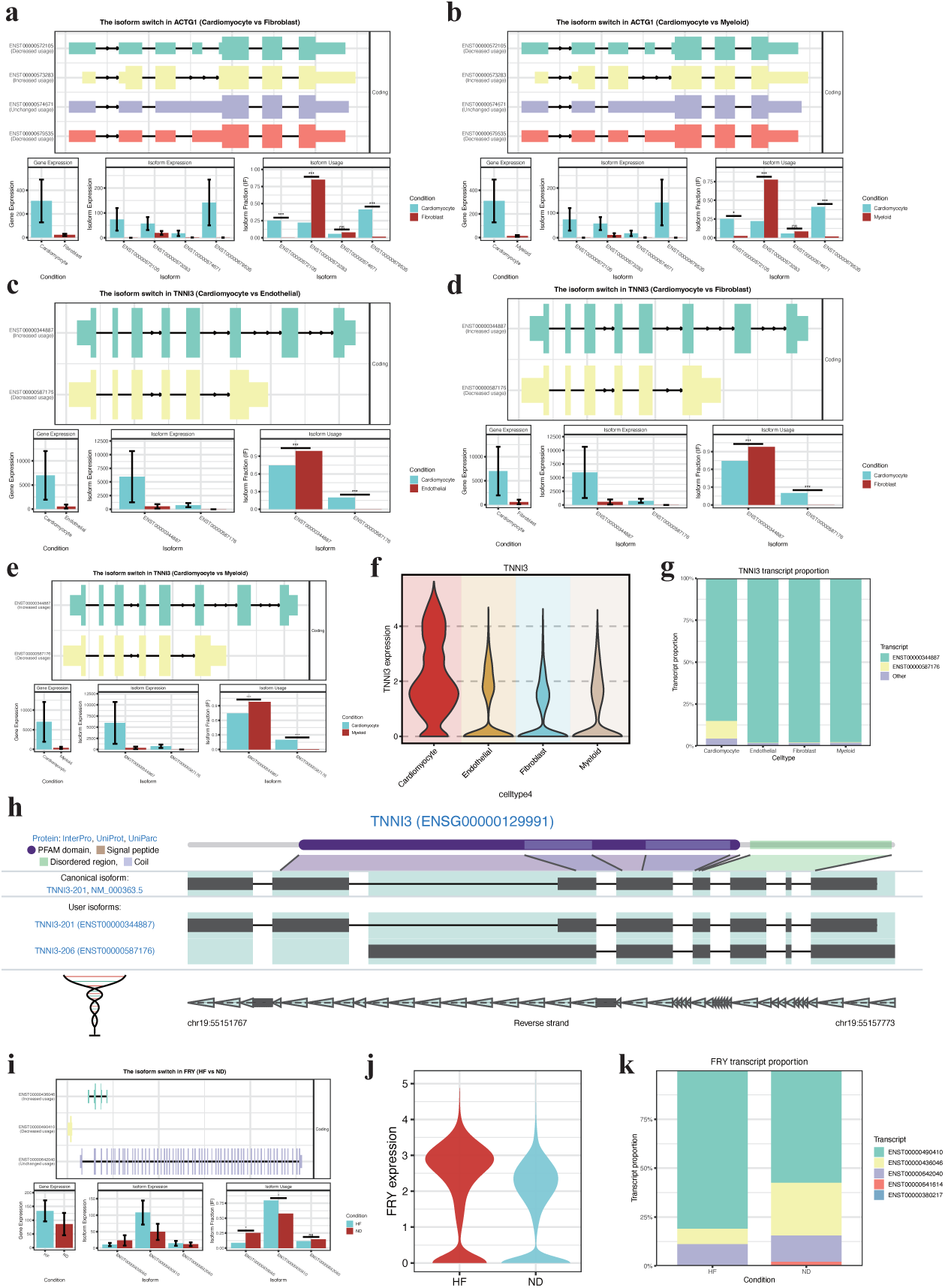
Representative differential transcript usage events reproduced by scIsoAgent. **a,b,** *ACTG1* DTU events from pairwise comparisons between cardiac cell types, with transcript structures and expression-based usage summaries. **c–h,** *TNNI3* DTU events from pairwise comparisons between cardiac cell types, with gene-level expression, transcript proportions, and transcript structure annotation. **i–k,** *FRY* disease-associated DTU event between heart failure and non-diseased cardiomyocytes. DTU, differential transcript usage; HF, heart failure; ND, non-diseased; ns, not significant; *, *p <* 0.05; ***, *p <* 0.001.

